# The Limits to Parapatric Speciation II: Strengthening a Preexisting Genetic Barrier to Gene Flow in Parapatry

**DOI:** 10.1101/266098

**Authors:** Alexandre Blanckaert, Joachim Hermisson

## Abstract

Parapatric speciation has recently received a lot of attention. By encompassing the whole continuum between allopatric and sympatric scenarios, it includes many potential scenarios for the evolution of new species. Building upon previous work, we investigate how a genetic barrier to gene flow, that relies on a single postzygotic genetic incompatibility, may further evolve. We consider a continent island model with three loci involved in pairwise Dobzhansky-Muller incompatibilities (DMIs). Using a deterministic and analytic approach, we derive the conditions for invasion of a new mutation and its consequences on an already existing genetic barrier to gene flow. We focus on quantifying the impact of the epistasis generated by the new mutation on the genetic barrier. We show that the accumulation of genetic incompatibilities in the presence of gene flow is a complex process, where new mutations can either strengthen or destroy a preexisting barrier. In particular, preexisting polymorphism and incompatibilities do not always facilitate the growth of the genetic barrier by accumulation of further barrier genes. Migration may disrupt the snowball effect (the accelerating rate of DMI accumulation in allopatry) because incompatibilities are directly tested by selection. Our results also show an ambiguous role of gene flow, which can either impede or facilitate the strengthening of the genetic barrier. Overall, our results illustrate how the inclusion of gene flow renders the building of a genetic barrier difficult to analyze.

## Introduction

Under what conditions can geographically separated populations that are connected by migration build up a genetic barrier to gene flow? When and how can this barrier be strengthened and eventually lead to speciation? Following the increasing awareness that gene flow and hybridization between related (incipient) species is ubiquitous in both plants and animals (Mallet, 2005; Butlin et al., 2008), these long-standing questions of parapatric speciation research are receiving renewed interest (Butlin et al., 2012; Bank et al., 2012; Flaxman et al., 2013, 2014; Paixão et al., 2014; Seehausen et al., 2014; Barnard-Kubow et al., 2016; Kulmuni and Westram, 2017; Nosil et al., 2017; Yang et al., 2017). Answers to these questions strongly depend on the speciation mechanism that is considered. On the one hand, there are scenarios of “adaptive speciation” (Dieckmann, 2004; Weissing et al., 2011), where speciation (or the build-up of a genetic barrier) is a direct target of selection. The genetic barrier in this case is usually prezygotic and can result from the evolution of assortative mating. If speciation is driven by local competition (as in the classical scenario of sympatric speciation, Dieckmann and Doebeli (1999)), the probability or speed of speciation is unaffected by migration. Alternatively, if assortative mating evolves as a response against mating with maladaptive immigrants, migration is driving speciation in the first place (Servedio and Noor, 2003; Rettelbach et al., 2013). On the other hand, other scenarios consider speciation as a non-selected by-product of neutral or adaptive divergence. In particular, this is how reproductive isolation evolves in classical models of allopatric speciation (Orr, 1995; Orr and Turelli, 2001; Coyne and Orr, 2004). In contrast to the scenarios of adaptive speciation, in that case, migration is a potent force to prevent the build-up of a genetic barrier. Given that models of adaptive speciation require specific assumptions about the selection scheme and given the ubiquitous nature of gene flow, the question arises whether and when speciation as a by-product can occur in a parapatric model.

Following previous work (Bank et al., 2012; Flaxman et al., 2013; Akerman and Bürger, 2014; Aeschbacher and Bürger, 2014; Paixão et al., 2014; Frässe et al., 2014; Höllinger and Hermisson, 2017), we study the conditions for the emergence of a postzygotic barrier to gene flow between parapatric populations. Two mechanisms can contribute to the build-up of such a barrier: local adaptation and genetic incompatibilities (Schluter, 2009; Bank et al., 2012; Kulmuni and Westram, 2017). Local adaptation and divergence driven by ecological differences among the populations is arguably the easiest mechanism to create a barrier in the presence of gene flow (Flaxman et al., 2013; Akerman and Bürger, 2014). Any new mutation with a local fitness advantage larger than the migration rate can establish in the population. If this same mutation is detrimental in the other environment, the adaptation remains local and contributes to a fitness deficit of migrants. An increasing number of local adaptation genes along the chromosome can strengthen the barrier and reduce the effective rates of gene flow among populations. Speciation in the sense of full reproductive isolation corresponds to the limit where immigrants are “dead on arrival”. However, hybridization remains possible whenever populations can overlap at all, in any environment (or laboratory) where both are viable. This problem is avoided if the genetic barrier is due to genetic incompatibilities and selection acts primarily on hybrids rather than on (first generation) migrants. This is the insight of the Bateson-Dobzhanszy-Muller model (Bateson, 1909; Dobzhansky, 1936; Muller, 1942) that has since become the standard model to explain speciation in an allopatric setting (Orr and Turelli, 2001; Coyne and Orr, 2004).

The two mechanisms, selection against migrants (i.e. local adaptation) and selection against hybrids (Dobshansky-Muller incompatibilities, DMIs) are non-exclusive (Kulmuni and Westram, 2017). In particular, whereas neutral DMIs cannot evolve in a parapatric setting (Gavrilets, 1997; Bank et al., 2012), DMIs can still evolve and be maintained if at least one of the incompatible alleles is also locally adaptive. Considering a continent-island scenario, Bank et al. (2012) characterized the conditions under which a simple 2-locus DMI can originate and be maintained in the face of gene flow – a very first step on the route to (potential) speciation. Here, we ask how this process can continue. Under which conditions will further substitutions in either population strengthen or weaken (or even destroy) an existing genetic barrier? It turns out that the answer to this question is surprisingly complex, depending on patterns of epistasis and on the genetic architecture and linkage pattern of the barrier genes involved. We discuss the potential of a new mutation to strengthen a barrier and whether it is a step towards reproductive isolation. Lastly, we characterize the genetic architecture that produced the strongest genetic barrier under gene flow and relate these results to the recent discussion of so-called “islands of divergence” (Via and West, 2008; Feder et al., 2012a).

## Model

To study the accumulation of incompatibilities in the presence of gene flow, we use a migration-selection model in continuous time with three loci. We consider two panmictic populations, one on a continent and the other on an island, each of sufficient size such that we can ignore the effects of genetic drift. There is unidirectional migration from the continental population to the island population at rate *m*. Selection acts on three loci, 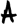, 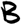, and 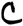, with two alleles each (**A/a, B/b, C/c**). Lower case letters indicate the ancestral state, upper case letters are derived alleles. We study both haploid and diploid populations. We always assume that the continent is fixed for a unique genotype; substitutions on the continent can occur, but they are instantaneous and do not lead to a persistent polymorphism. We focus on the migration-selection dynamics on the island, where all three loci can be polymorphic.

## Haploid model

There are 2^3^ = 8 different haplotypes with frequencies *x*_1_, *x*_2_, …*x*_8_. In particular, *x*_1_ is the frequency of the ancestral genotype **abc**, with Malthusian (or log-) fitness normalized to 0. We have three parameters for single-locus fitness effects, *α*, *β*, and *γ*. Three parameters *∊_AB_*, *∊_BC_*, and *∊_AC_*, parametrize potential pairwise epistasis between derived alleles, see table 1. Restrictions on epistasis values are detailed below.

**Table 1:** **Frequencies** *x_i_* **and fitness values** *w_i_* **of the different haplotypes for haploid populations. We always assume** *α* > 0 **and** *∊_AB_* < 0.

| Hap. | **abc** | **Abc** | **aBc** | **abC** | **ABc** | **AbC** | **aBC** | **ABC** |
| --- | --- | --- | --- | --- | --- | --- | --- | --- |
| $x_i$ | $x_1$ | $x_2$ | $x_3$ | $x_4$ | $x_5$ | $x_6$ | $x_7$ | $x_8$ |
| $w_i$ | 0 | $\alpha$ | $\beta$ | $\gamma$ | $\alpha + \beta$<br>$+ \epsilon_{AB}$ | $\alpha + \gamma$<br>$+ \epsilon_{AC}$ | $\beta + \gamma$<br>$+ \epsilon_{BC}$ | $\alpha + \beta + \gamma$<br>$+ \epsilon_{AB} + \epsilon_{BC} + \epsilon_{AC}$ |

In the following, we assume that each locus has a specific role. In particular, we assume that allele **A** is always an island adaptation (allele **A** appears on the island). As a consequence, α is always strictly positive. In contrast, allele **B** is always a continental adaptation (allele **B** appears on the continent). There is no constraint on its selective advantage, *β*, on the island: both negative and positive values are investigated. We always assume that **A** and **B** are incompatible, i.e. *∊_AB_* < 0. While loci 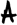 and 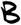 form the nucleus of a genetic barrier that exists initially, any further extension of this barrier occurs on the 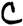 locus. At this locus, the new allele **C** can appear either on the island or on the continent. There is no constraint on its selective advantage, *γ*. **C** can interact positively or negatively with the other derived alleles. To keep our model tractable, we only allow for epistasis between island and continental adaptations. In other words, if **C** appears on the island, it only interacts with the continental adaptation **B** (*∊_AC_* = 0). Similarly, if **C** appears on the continent, epistasis only occurs between **A** and **C** (*∊_BC_* = 0). This excludes schemes of complex epistasis with interactions among all three locus pairs, or higher-order interactions.

Note that our choice for the role of loci 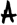, 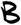, and 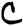 is made to reduce the parameter space. Alternative scenarios can be easily deduced through reparametrization of the system, given in table A3 in the SI. Since the model is defined in continuous time, all parameters for selection or migration are rates. For the derivation of equilibria, only relative rates matter. In particular, we can scale all parameters by the selection coefficient of the **A** allele (which is always > 0).

The three loci 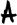, 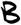 and 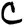 can be located in any order along the genome. The full system with arbitrary linkage, given in equation (A2) is not tractable analytically. In our analysis, we therefore focus on limiting cases with pairs of loci either in tight linkage (recombination rate *r* → 0) or in loose linkage. In our model, we implement loose linkage as the limit *r* → ∞, which implies that the corresponding loci are always in linkage equilibrium. We relax this assumption in the SI, Fig. C24,C25, where we discuss numerical results for the dynamics with intermediate recombination. The linkage equilibrium approximation holds as soon as recombination is stronger than the other evolutionary forces (selection and migration). This gives rise to five different linkage architectures: 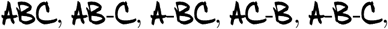 where “-” denotes loose linkage and its absence tight linkage. We investigate these architectures both for 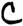 appearing on the island or on the continent respectively, leading to 10 different cases.

The dynamical equations for the allele frequencies on the island (*p_A_*, *p_B_*, *p_C_* for allele **A**, **B**, **C**, resp.) for all cases are derived in the SI, equations (A6)-(A8). For example, we obtain for loose linkage 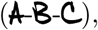

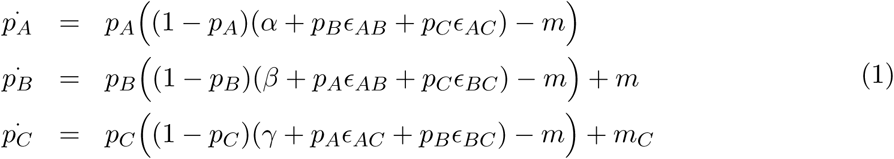

where *m_C_* = *m* or *m_C_* = 0, depending on whether **C** appears on the continent or on the island.

## Diploid model

We define the fitness scheme for diploids as follows: single-locus effects (*i.e. α*, *β*, *γ*) are purely additive. There is thus no dominance at this level. Dominance is, however, included for epistasis. Following previous work (Turelli and Orr, 2000; Bank et al., 2012), we assume that the strength of epistasis depends only on the number of incompatible pairs in a genotype, e.g. **AB/Ab** generates the same epistasis as **AB/aB**.

We investigate two cases of dominance of the epistatic interaction: recessive and codominant epistasis (see table 2). Assuming Hardy-Weinberg equilibrium on the island, the dynamical system for diploids coincides with the haploid equation (given in equation (A2)) if we replace the fitness of all haplotypes by the corresponding marginal fitness 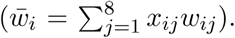 In the case of the codominant model, the diploid dynamics reduce to the dynamics of the haploid model if all interacting loci are in loose linkage, 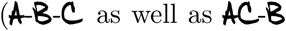 if **C** appears on the island, and 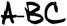 if **C** appears on the continent). The different systems of equations are available in the SI, (see equations (A11)-(A15)).

**Table 2:**
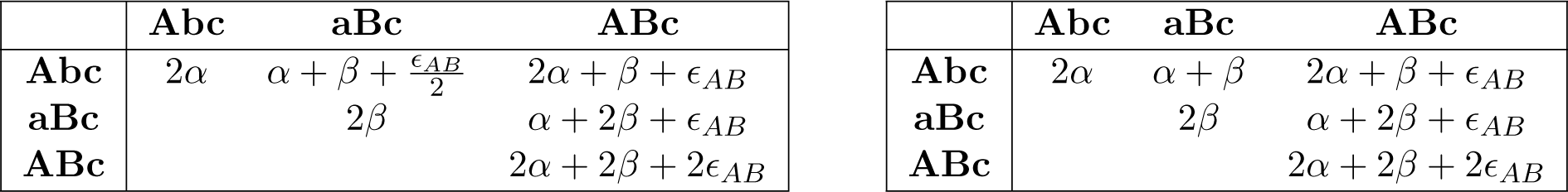
**Section of the fitness table specifying the interactions between the A and B alleles in the background of allele c for codominant (left) and recessive (right) epistasis. Interactions between A and C as well as B and C are analogous (the complete table is available in the SI, table A2).**

## Strength of the genetic barrier

There are multiple measures for the strength of a genetic barrier between two divergent populations that are connected by gene flow. For example, the gene-flow factor (or the effective migration rate) due to Barton and Bengtsson (1986) measures the reduced probability of neutral alleles that are linked to barrier genes to cross this barrier and establish in the recipient population. Here we consider the fate of barrier genes themselves. In particular, we are interested in the maximum rate of gene flow under which a barrier (with given selection parameters) can be built and also in the maximum rate of gene flow under which such a barrier can persist if it exists initially.

Specifically, we define the barrier strength 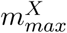 for a given set of barrier loci as the maximal migration rate under which a set X of alleles at these loci can still be maintained on the island. Here, X denotes the barrier alleles that are not present on the continent, but are maintained on the island as long as migration is below the threshold 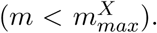 For example, for a single-locus barrier with the **A** allele on the island, we have X = **A** and the strength of the genetic barrier is given by 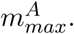 For 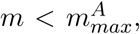 the 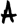 locus is polymorphic on the island, for 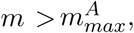 the **A** allele is swamped and the locus is fixed for the continental **a** allele. Analogously, the strength of a genetic barrier with three polymorphic loci and island alleles **A**, **b**, and **C** (say) is denoted as 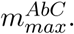 The two-locus barrier 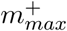 from Bank et al. (2012) corresponds to 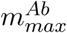 with this notation.

Below, we consider how the strength of an existing genetic barrier changes under further evolution. We then denote the original barrier strength, which serves as the reference point, as 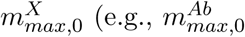 is the initial strength of an 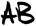 barrier with the third locus 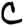 fixed for its ancestral allele **c**). While 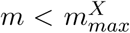 guarantees that an existing DMI is not swamped, the origin of the DMI may require a favorable evolutionary history (mutation order) or an initial allopatric phase, (Bank et al., 2012).

## Results

### Adaptation at existing barrier loci

In the first part of the results section, we study the case where further adaptation happens directly at an already existing barrier locus (i.e., at a locus 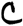 in tight linkage to such a locus). In particular, we compare the simple case of further adaptation at a single barrier locus with the more complex scenario where adaptation happens at a barrier locus that is involved in a 2-locus incompatibility.

#### Further adaptation at a single-locus barrier

Assume that, initially, 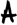 is the only polymorphic locus. The initial barrier strength is *α* and results entirely from local adaptation 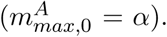 A new mutation occurs at a tightly linked locus, 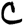. This scenario is equivalent to adaptation at a single compound locus 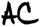 with alleles **Ac** and **ac** and mutation generating new alleles **AC** with fitness *α* + *γ* and **aC** with fitness *γ*. At most two alleles can be maintained on the island (Nagylaki and Lou, 2001): the continental allele and the allele with the highest fitness on the island. Thus, any new adaptation on the island that produces a better allele than **Ac** will replace this allele (eg., the **AC** allele for *γ* > 0). While any successful adaptation on the island increases the barrier strength 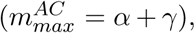 adaptation on the continent can lead to a stronger or weaker barrier 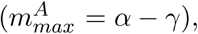 depending on whether *γ* is positive or negative. In particular, 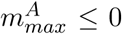 means that no polymorphism can be maintained. However, if there is a small but non-zero recombination probability among 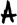 and 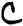, any adaptation on the continent will eventually also enter the island background. We then have two new alleles **AC** and **aC** replacing the old ones (**Ac** and **ac**) and the barrier strength 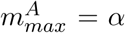 remains unchanged as long as there is no epistasis among **A** and **C**.

We thus see that further adaptation at a single polymorphic locus will usually strengthen the genetic barrier, rather than weaken it. In particular, this holds for any further adaptation on the island. The only exception is adaptation on the continent that is also beneficial on the island and cannot be combined with an existing island adaptation (i.e. the combined allele **AC** is not possible or deleterious). A 3-locus architecture 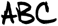 with tight linkage among all three loci leads to an analogous single-locus problem (after appropriate relabeling of parameters).

Note that the genetic barrier formed by a single locus relies exclusively on local adaptation: any isolation observed is due to the impossibility of coexisting in a common environment and not due to a genetic mechanism. This is different for barriers with multiple interacting loci, which is our focus in the remainder of the manuscript.

#### Further adaptation at a two-locus barrier

Assume now that we start with a 2-locus polymorphism at two incompatible loci 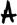 and 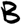 (a 2-locus DMI) in loose linkage. The continental haplotype is **aB**, and **Ab** is the fittest haplotype on the island. A new mutation appears on the island at a locus 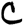 in tight linkage with 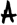. As discussed in the previous section, this generates a compound locus 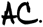 The new mutation generates a third allele at this compound locus (e.g. **AC**), which we will call the **A’** allele in the following. We denote the fitness advantage of the new allele **A’** as *α^′^* and its epistatic interaction with the **B** allele at the 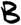 locus as *∊_A′B_*. This leads to the dynamics of a triallellic locus (with alleles **a**, **A** and **A’**) that interacts with a loosely linked biallelic locus in the genomic background (alleles **b** and **B**):

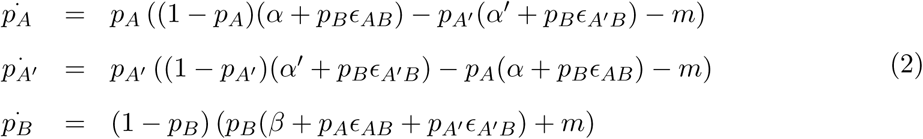

(For tight linkage, we can assume that a fourth allele **A”** (e.g. **A”** = **aC**) will only originate by mutation or rare recombination after one of the alleles **a**, **A**, and **A’** is lost. This leads again to the three-allele dynamics described by Eq. (2). Results for the four-allele dynamics are given in the SI, section C 2.4)

The dynamical system, given in equation (2) allows up to 9 equilibria, up to 3 of which can be simultaneously stable. In the SI, section B2, we show that alleles **A** and **A’** can never coexist at a stable equilibrium (extending the single-locus result of Nagylaki and Lou (2001)). Nevertheless, interaction of 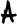 with an unlinked locus 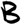 considerably adds to the complexity and can lead to qualitatively different results.

Whereas **A** and **A’** can not coexist, allele **A’** can still invade the equilibrium formed by the DMI between loci 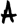 and 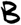. In contrast to the single-locus case, the potential for **A’** to invade does no longer depend only on the fitness values, but also on the strength of migration (analytical expressions of the bounds are given in equation B7). In Fig. 1, invasion of **A’** is possible in all colored regions. Fig. 1(a) shows invasion of an allele **A’** with larger direct effect *α^′^* > *α*. If negative epistasis is less severe for **A’** than for **A** (*∊_AB_* < *∊_A′B_* < 0), **A’** will always invade (i.e. up to the maximal migration rate, 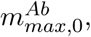 of the original two-locus polymorphism). However, for strong negative epistasis of the new allele (*∊_A′B_* < *∊_AB_* < 0) invasion of **A’** is only possible for weak migration 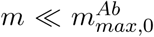 and a sufficiently low frequency of the competing **B** allele on the island. Fig. 1(b) shows that the **A’** allele can also invade if its direct effect is weaker (*α^′^* < *α*), provided negative epistasis is also weaker (*∊_AB_* < *∊_A′B_* < 0). This requires that migration is sufficiently strong, because the marginal fitness of **A’** becomes larger than the marginal fitness of **A** only for a sufficiently large frequency of **B** alleles.

**Figure 1:**
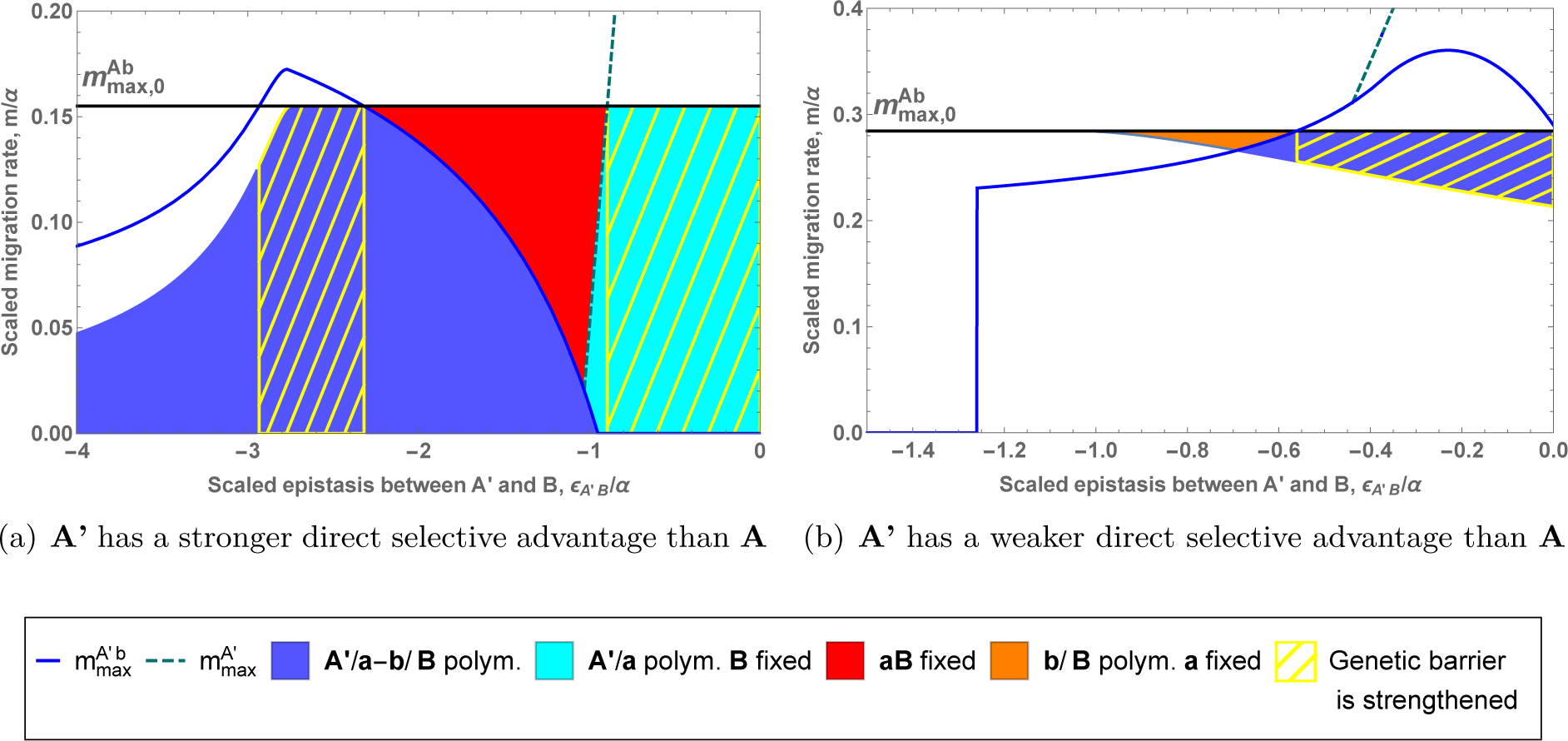
The impact of the invasion of a new allele A’ at locus 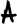 on an existing 2-locus barrier (loci 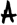, 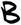 in loose linkage). We show the strength of a genetic barrier to swamping for a new allele **A’** as a function of its (scaled) epistatic coefficient. The strength of the original genetic barrier is indicated by the black line. In both examples, we have 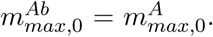 Invasion of allele **A’** can only happen in a finite interval for *m*, (equation (B7) for explicit expressions), corresponding to the colored area. There are four possible outcomes to the successful invasion of the **A’** allele, denoted by the background color: **A’** replaces **A** and **b** remains present (in blue), **A’** replaces **A** but allele **B** fixes (in cyan), the polymorphism at locus 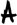 is lost (orange) and the continental haplotype fixes (red). If **A’** successfully replaces **A**, the new 2-locus barrier strength, 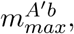 is given by the blue line. The dashed cyan line shows the strength of the new single-locus barrier, 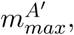 whenever the 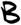 locus is swamped and 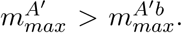 The yellow hatched area indicates that the genetic barrier at the 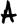 locus is strengthened by the invasion of allele **A’**. Panel a) is obtained for 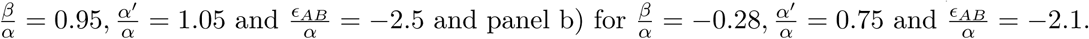

Successful invasion of **A’** can have qualitatively different outcomes, indicated by the different colors in Fig. 1. In many cases, an invading **A’** allele displaces the old **A** allele and the system settles at a new equilibrium with an **a**/**A’** polymorphism. The new equilibrium can either be a two-locus polymorphism (blue areas in Fig. 1) or a single-locus polymorphism with the B locus fixed for the **B** allele (cyan area). In both cases, the strength of the genetic barrier with respect to swamping can either increase (blue or cyan line above the black line) or decrease (blue or cyan line below the black line). Parameter ranges where invasion leads to a stronger genetic barrier, 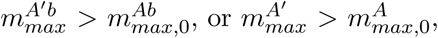 are indicated by yellow hatches.

Strengthening of the 2-locus barrier (blue area with yellow hatches in Fig. 1) can be due to two mechanisms. First, selection against migrants can be stronger due to additional local adaptation (*α*^′^ > *α*) and therefore leads to a larger fitness deficit (*α^′^* − *β*) for the continental haplotype on the island. This is the same mechanism as for the single-locus case. The genetic barrier is strengthened as long as epistasis, *∊_A′B_*, does not deviate too much from the epistasis generated by the previous allele, *∊_AB_* (Fig. 1(a), blue line above the black line). Indeed, if epistasis is too weak, the boost provided by the increased selection against migrants is negated by the weakening of selection against hybrids (since *β* > 0). If epistasis is too strong, on the other hand, the marginal fitness of allele **A’** is decreased due to the increased cost of hybrids. Allele **A** can invade such an equilibrium as soon as migration increases and **A’** cannot strengthen the genetic barrier (see also section B 6 in SI).

The alternative mechanism corresponds to the reduction of selection against hybrids, Fig. 1(b). It works only if the continental **B** allele is deleterious on the island. Indeed, in this scenario, selection against hybrids does not contribute to the genetic barrier, as **B** is already maladaptive on the island. Nevertheless, epistasis still generates a cost for the island adaptation through the production of hybrids. Therefore, releasing the selective pressure on locus 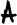 due to the hybrid cost (*∊_AB_* ≪ *∊_A′B_*) can strengthen the genetic barrier, even if this relief is associated with a reduction of the direct selective advantage of the island adaptation (*α^′^* < *α*). The reduction of the selection against migrants is here compensated by the much lower hybrid cost paid by allele **A’** relative to allele **A**.

In contrast to the single-locus case, invasion of **A’** does not imply that this allele is maintained in the population. Indeed, we find significant parameter regions, where the following scenario happens. First, allele **A’** invades the island population (at its initial equilibrium with two-locus polymorphism), leading to the loss of the **A** allele. In the absence of allele **A**, allele **B** is no longer repressed and increases in frequency, making it impossible for allele **A’** to maintain itself in the population. Consequently, the continental **a** allele swamps the island and the polymorphism at the 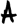 locus is lost altogether (red and orange areas in Fig. 1). Again, the polymorphism at the 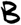 locus can either be maintained (orange area, Figure 1(b)) or destroyed (red, Fig. 1(a)). Clearly, a necessary condition for such behavior is that the original 2-locus polymorphism is not globally stable in the original **a**/**A**, **b**/**B** state space, but bistable together with an equilibrium with the **a** allele fixed. Numerical evidence strongly suggests that the fate of an invading **A’** allele depends on the existence of a stable **a/A’** polymorphism in the state space spanned by **a**/**A’**, **b**/**B**. If it does, the **A’** allele will eventually establish (as discussed above), if it does not, the **a** allele will take over. (We did not find a case where the **A** allele would return and displace **A’** once the latter has been able to invade; see also section B 3 in SI, for a more detailed discussion and some proofs for specific cases).

A new allele **A’** can thus function as a temporary state that enables switching among different equilibria of the original 2-locus 2-allele system. In the examples discussed above, temporary invasion of **A’** will destroy a DMI polymorphism in this case. However, we also observe the opposite phenomenon: invasion of **A’** may create an **a**/**A**-**b**/**B** polymorphism rather than destroying it. This is illustrated in Fig. 2. Bank et al. (2012) described the origin of such a DMI as a result of secondary contact (or a similar starting condition). Here, we provide an alternative explanation that does not require an interruption of gene flow.

**Figure 2:**
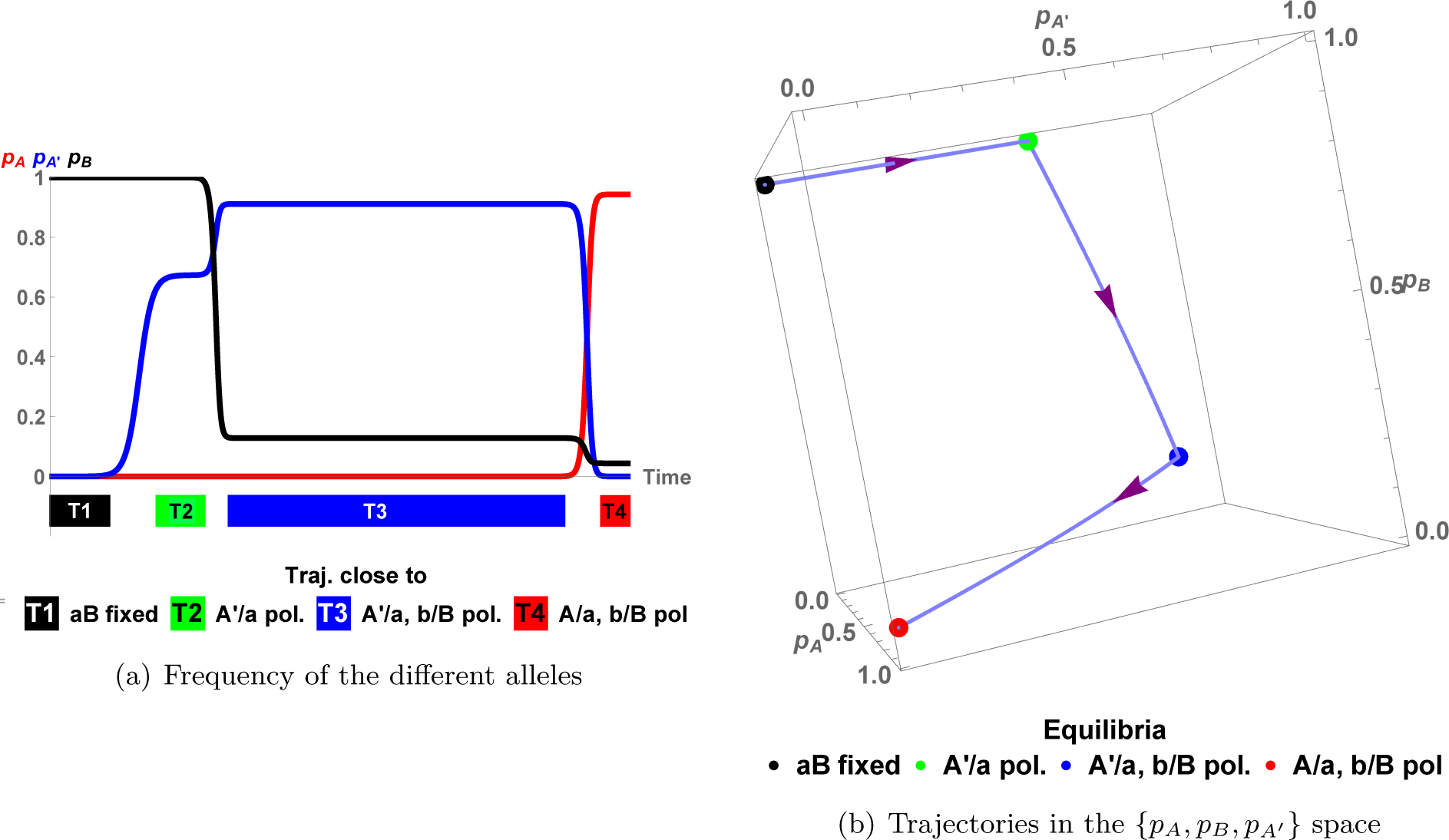
Evolutionary trajectory with A’ as transient state. a) Frequency of derived alleles **B** (black), **A** (red), and **A’** (blue) as a function of time. At t=0, the population is almost monomorphic, for the continental haplotype **aB**, with both the allele **A** and **A’** present at an extremely low frequency (≈ 10^−6^). Colored blocks T1-T4 indicate when the population is close to an equilibrium, with the color matching the corresponding equilibrium. b) We represent the same trajectory in the {*p_A_*, *p_B_*, *p_A′_*} space. Dots indicate equilibria, as indicated. Arrows indicate the evolutionary trajectory. Parameters used are: 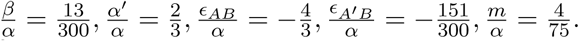 One can observe a similar behavior with locus B starting polymorphic and allele **B** deleterious on the island, see Fig. B2.

We can compare the consequences of further adaptation on the island at an existing genetic barrier in the two cases discussed so far: a single polymorphic locus 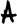 and a polymorphic locus 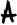 that interacts with a second polymorphic locus 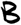. There are two notable differences:

*•* While further adaptation on the island always leads to a stronger barrier in the single-locus case, this is not the case for a 2-locus barrier. Furthermore, invasion of a new allele no longer even guarantees establishment of this allele. On the contrary, we see that such an event can erase the existing barrier entirely.

*•* The potential to strengthen the genetic barrier does not only depend on the fitness landscape, but also on the migration rate. Suppose that an allele **A’** exists that leads to a stronger barrier than **A** – if it invades. Fig. 1 shows that invasion may either require sufficiently weak (Fig. 1(a)), or sufficiently strong migration (Fig. 1(b)). The latter scenario leads to the interesting observation that stronger gene flow can sometimes trigger the evolution of stronger barriers to gene flow (in Fig. 1(b), 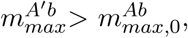 the blue line is above the black line. Invasion of the new mutant is only possible with relatively strong migration (colored area in the figure)).

We also observe a general trend to replace a polymorphism that is maintained by selection against hybrids by one that is maintained due to selection against immigrants. Indeed, whereas it is possible to strengthen the genetic barrier by weakening the strength of epistasis without affecting the amount of local adaptation, the opposite is impossible. Any increase in the strength of selection against hybrids needs to be associated to some increase in local adaptation.

Further adaptation at locus 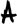 on the continent or at locus 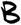 on the island are equivalent to the one locus case, since the new alleles, **a’** or **b’** cannot generate epistasis with **B** or **A** respectively (assumption of the model). Further adaptation at 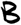 on the continent is treated in the SI, section B7. If *β^′^* < 0, strengthening of the barrier is more likely with weaker epistasis, whereas if *β^′^* > 0 most of strengthening happens if the incompatibility gets stronger. If **B’** is much more deleterious than **B** on the island, the genetic barrier is strengthened regardless of epistasis.

### Extension of the genetic barrier

We now turn to the extension of a genetic barrier by adaptation at an interacting locus 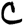 that is far away from the existing barrier loci and only loosely linked. We start with going from one to two loci and then study the case when a third locus is added.

#### Extension of a single-locus genetic barrier

Assume that 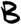 is the only polymorphic locus on the island 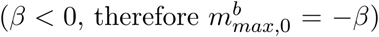 and a new mutation **C** occurs on the island at a loosely linked locus 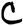. In the absence of epistasis, this mutation can invade and establish if and only if *γ* > *m*. 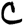 does not affect the barrier at all.

Fig. 3(b) shows the effect of epistasis between **C** and **B** on the barrier strength. As expected, negative epistasis can strengthen the genetic barrier (blue area), while positive epistasis will almost always weaken it (orange and red areas). The figure also shows, however, that negative epistasis is not sufficient to strengthen the barrier. Indeed, a **C** allele may invade for sufficiently weak migration, but will be the first polymorphism swamped once migration increases (grey area). Obviously, in this case, the barrier strength 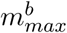 at the polymorphic 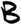 locus remains unaffected.

**Figure 3:**
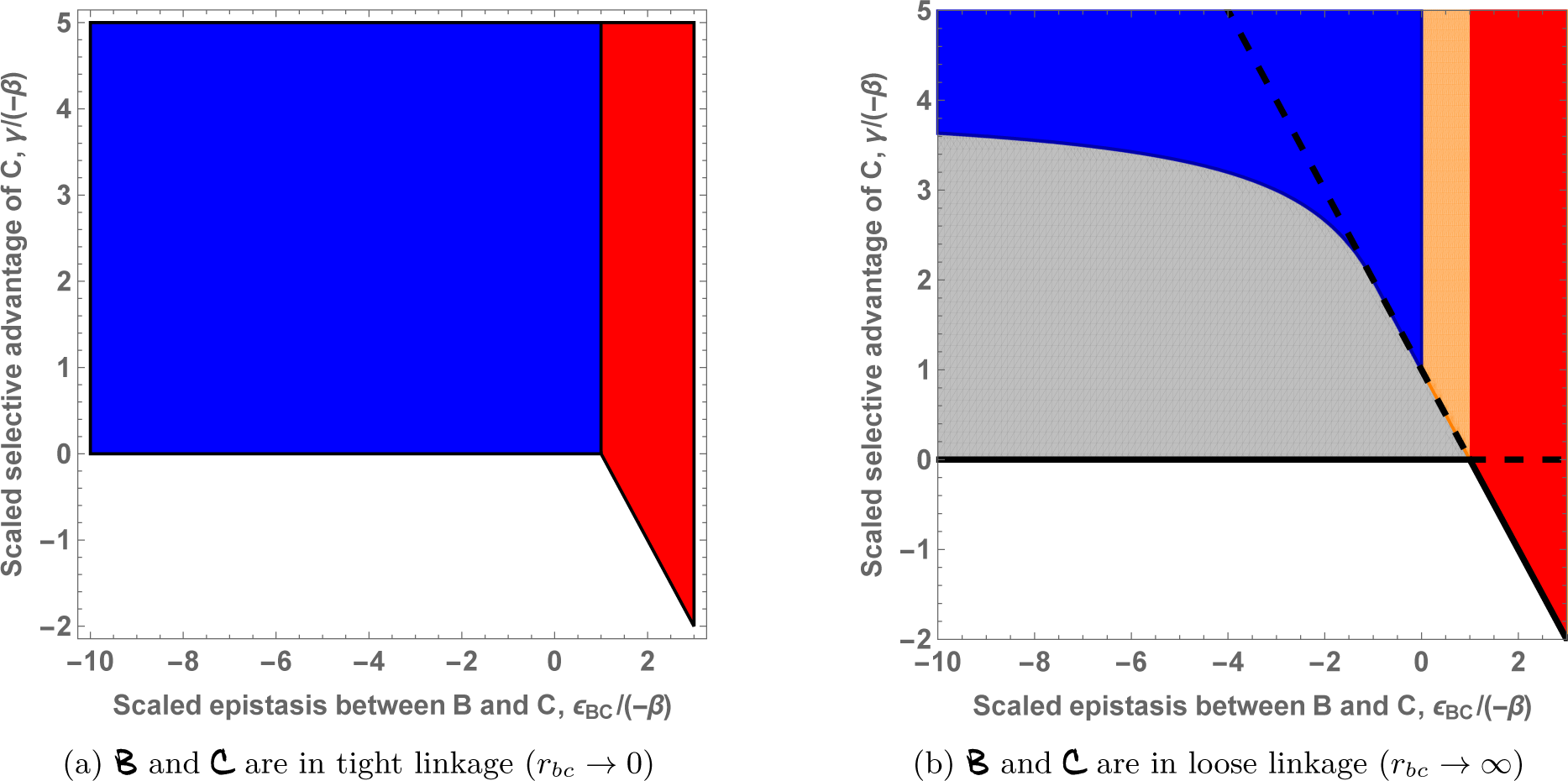
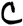 strengthens the genetic barrier formed by a single polymorphic locus: comparison between a new mutation in tight linkage and one in loose linkage. The x-axis shows the strength of epistasis between 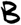 and 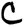. The y-axis shows the selective advantage of new allele **C**. The background color indicates the consequence of the invasion of allele **C** on the genetic barrier at the 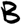 locus. Gray: the genetic barrier remains unchanged; blue: the genetic barrier is strengthened; orange: the genetic barrier is weakened; red: the polymorphism at locus 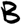 is lost. In addition, on panel b), the solid black line gives the necessary condition for invasion of allele **C** on the island. Below this bound invasion is always impossible. The black dashed line gives the sufficient condition for invasion. Above this bound, allele **C** can always invade, regardless of the migration rate (provided the polymorphism at the 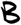 locus still exists). Analytical expressions for the two lines are given in SI, equation (C2).

It is instructive to see how linkage affects the parameter range where further adaptation leads to a stronger barrier. For tight linkage (adaptation at the polymorphic locus itself), any allele with *γ* > 0 will invade the island and will strengthen the barrier, see Fig. 3(a), as long as epistasis does not cancel the selective disadvantage of allele **B** (*∊_BC_* + *β* > 0). In contrast, strengthening the barrier by adaptation at a loosely linked locus is much more difficult. To reinforce the barrier at the loosely linked 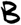 locus, the new **C** allele has to withstand both migration for 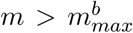 and the hybrid cost generated by its interaction with allele **B**. The first condition alone implies *γ* > −*β* as a necessary condition for a stronger barrier. Indeed, for a given *γ* > −*β*, the genetic barrier 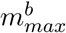 is strengthened as long as negative epistasis is not too strong. Stronger epistasis results in larger hybrid cost for **C** and therefore a larger direct effect (larger *γ*) is needed to compensate for it. For *γ* > −4 *β*, the barrier is strengthened for any negative epistasis, including a lethal incompatibility.

Finally even if a new **C** allele would strengthen the barrier (blue area), it is not always able to invade. Invasion of allele **C** requires

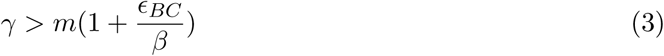

From equation (3), one can deduce that a necessary condition for invasion of allele **C** is *γ* > 0 and a sufficient one is *γ* > − (*β* + *∊_BC_*). For any *γ* value between these two limits, invasion will be possible only if migration is sufficiently small. Such a constraint does not exist for the tight linkage case, as migration does not affect the fate of new allele (given 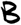 remains polymorphic).

So far, we have considered the interaction of a new island adaptation **C** with a continental adaptation **B**. Alternatively, there are two others possibilities: we can also study interactions between **B** and a new continental adaptation or interactions among two island adaptations. The results are similar, see Fig. C2, C3 and our discussion in the SI.

#### Extension of a two-locus genetic barrier

To complete the analysis of this section, we now ask how an existing genetic barrier of two interacting loci in loose linkage is affected by adaptation at a third locus that is also in loose linkage with the previous ones. We focus, in particular, on the question how a continental allele (the **B** allele at the 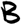 locus) can be prevented from swamping the island. Depending on the direct fitness effect *β* of this allele on the island, we find similarities or differences to the extension from 1 to 2 loci discussed above.

If the continental adaptation **B** is deleterious on the island (*β* < 0), direct selection against migrants (all carrying allele **B**) contributes to the genetic barrier, 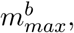 at that locus. As Fig. 4(a) shows, transition from two to three loci is analogous to the step from 1 to 2 loci and also the qualitative results agree (see SI section C 1.2 for details). Indeed, the presence of a first island adaptation (the **A** allele) does not make it any easier for a second, loosely linked island adaptation (the **C** allele) to strengthen the genetic barrier. In particular, the **C** allele still needs to have a stronger direct effect (in magnitude) than **B**, *γ* > −*β*. Allele **C** also needs to interact negatively with allele **B**, *∊_BC_* < 0. Finally, since this interaction generates some hybrid cost, this cost must be compensated by some extra local adaptation (larger *γ*). For example, a new mutation, interacting with allele **B**, with a direct selective advantage *γ* slightly larger than −*β*, might not be able to strengthen the genetic barrier even if it fulfills the first criteria. These conditions are analogous to the 1 to 2-locus barrier transition, see Fig. 3(b), gray area above the 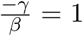 line. However, **C** can be weaker than the **A** adaptation (*γ* < *α*) and still lead to a stronger barrier, see Fig. 4(a).

**Figure 4:**
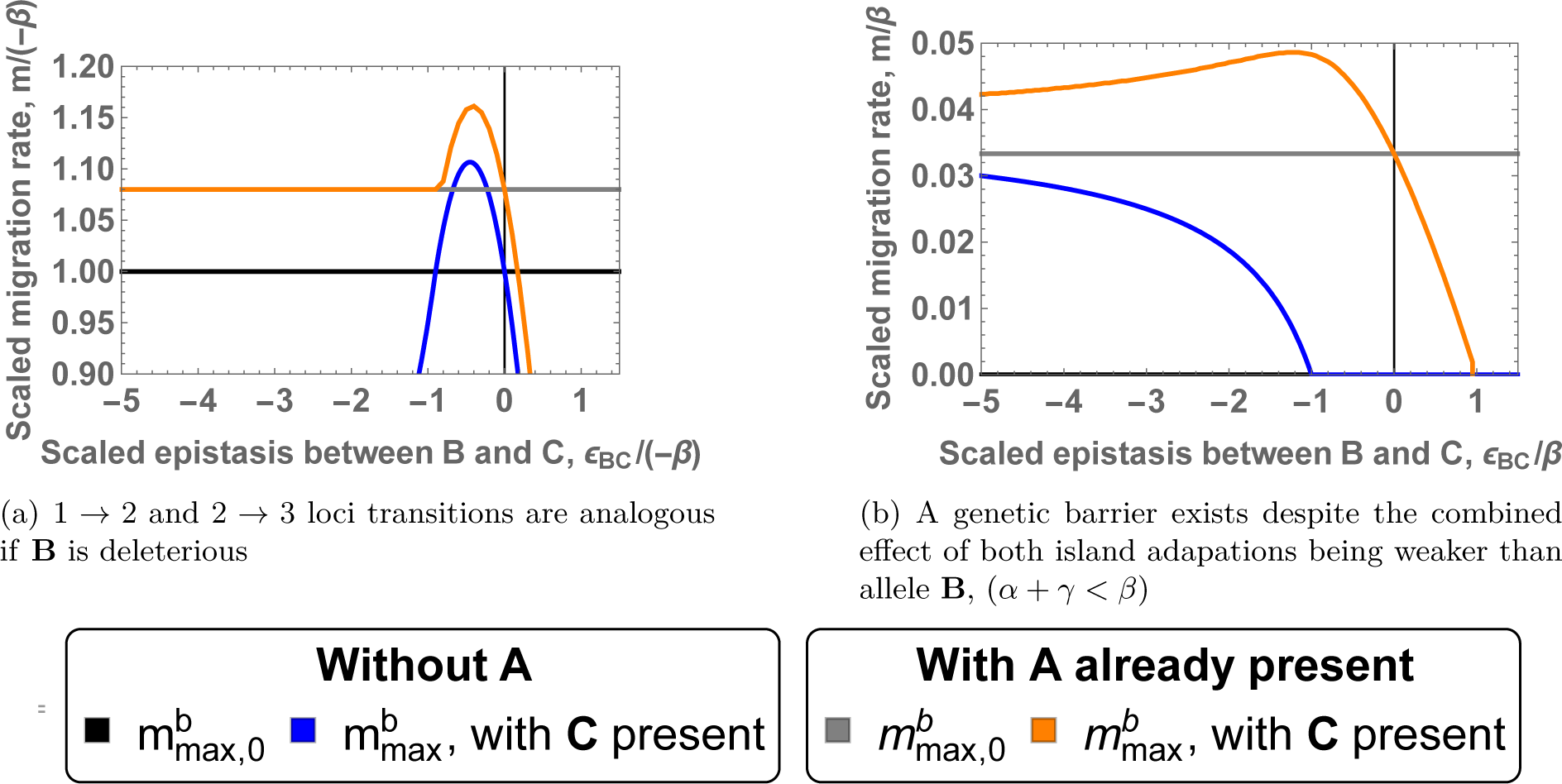
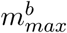 for two and three loci in loose linkage. The x-axis corresponds to the epistasis between alleles **B** and **C**. The y-axis measures migration rate. The different lines correspond to 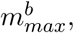 the resistance to swamping at locus 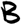 under different scenarios. The initial single-locus and two-locus barriers, 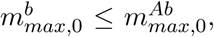 are given in black and gray. The impact of a new allele **C** on the single-locus and two-locus genetic barrier is represented by the blue and orange lines, respectively. Allele **B** is deleterious on the island for panel a) and advantageous on the island for panel b). The thin vertical black line indicates the absence of epistasis. Panel a) is obtained for 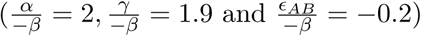 and panel 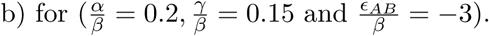

We now consider a continental allele **B** that is beneficial also on the island, Fig. 4(b). In the haploid model, a single-locus genetic barrier is impossible. A genetic barrier can be formed if a second polymorphic locus, 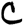, interacts with 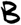 through negative epistasis, generating selection against hybrids. However, a stable genetic barrier only exists if the direct and the epistatic effect of the **C** allele are both strong, *∊_BC_* < −*β* − *m* and *γ* > 4*m*, (see section C 1.1.2 in the SI for details), represented by the blue line in Fig. 4.

Consider now such a two-locus barrier between loci 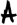 and 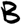. We want to investigate under which conditions a polymorphism at a loosely linked locus C strengthens the barrier against swamping at the 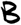 locus, 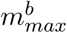 (orange line on Fig. 4). With an **A** allele already present, there is no lower bound for the negative epistasis of the new **C** allele: any value *∊_BC_* (*∊_BC_* < 0) can increase the barrier strength. The new allele still has to fulfill a condition on the direct effect: 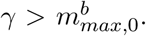 Otherwise, allele **C** is the first allele that is lost when gene flow increases. However, the condition is weaker than the one on the **A** allele; indeed *γ* > *α*/4 is a sufficient condition. Allele **C** can even have the weakest direct effect (Fig. 4(a)) and still contribute to the strengthening of the barrier, in contrast to the case *β* < 0 discussed above. The two island adaptations share the cost of forming hybrids, making it possible to prevent a strongly advantageous continental allele to fix on the island, despite their own relatively weak selective advantage (*α* + *γ* < *β*) (Fig. 4(a)).

Not only the maintenance, but also the invasion of the new polymorphism in loose linkage is strongly affected by the existence of a polymorphism at locus 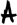. In its absence, the new mutation has to overcome the migration cost and the full incompatibility due to **B** being already fixed *γ* > *m*−*∊_BC_*. In addition, epistasis has an ambiguous effect: it hinders invasion of the new allele while the formation of the 2-locus genetic barrier requires relatively strong negative epistasis (*∊_BC_* < − *β* − *m*). This ambiguous effect makes invasion and establishment of a genetic barrier in this setting extremely unlikely. Once a two-locus genetic barrier exists, invasion of a new allele is however much easier and always possible if migration is sufficiently small (the invasion criterion tends to *γ* as *m* → 0). Invasion of a third mutation is therefore similar to the previous case (allele **B** is deleterious on the island).

As we have seen, the constraints on the **C** allele are not so severe - but the flip side is that also the effect on the barrier strength is quite weak: roughly 10% of the direct effect of the **C** allele for Fig. 4(a) and 5% for Fig. 4(b). In comparison, when all loci are in tight linkage, 100% of the direct effect of the new mutation contributes to strengthening the genetic barrier.

In the supplement, we explore a slightly different scenario, where the new mutation **C** does not interact directly with allele **B** but with allele **a**. Our results show that indirect strengthening of the genetic barrier, by increasing the marginal fitness of the **A** allele, can be the most efficient scenario (Fig. C6).

#### Barrier strength and linkage architecture

Assume now that 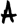, 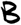, and 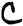 are placed without restrictions on recombination distance. For a given set of selection parameters, which linkage architecture will form the strongest barrier?

For a two-locus genetic barrier (loci 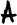 and 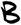), this question has been addressed by Bank et al. (2012). The main finding there is that selection against migrants is strongest for tight linkage while selection against hybrids is maximal in loose linkage, when most incompatible hybrids are produced. With both factors acting, the strongest barrier still results from one of these extreme architectures: 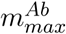 is maximized for tight linkage whenever selection against migrants is the main driving force. This is the case, in particular, whenever **B** is deleterious on the island (Fig. C20(a)). In contrast, selection against hybrids is the only viable factor if the continental type, **aB**, has the highest fitness also on the island. In this case, we obtain the maximal 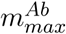 in loose linkage (Fig. C20(b)). Assuming a genetic barrier can be formed both in tight and loose linkage, the loose linkage architecture forms the strongest barrier if:

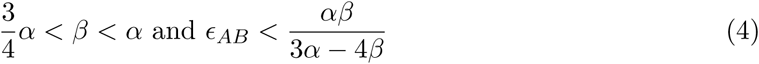

In particular, there is never a maximum for intermediate recombination rates.

The case of three loci is more complicated because conflicting options can exist, e.g. the strongest barrier for pairs 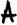 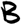 and 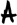 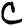 is obtained with the different loci in tight linkage, but the strongest barrier for the pair 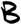 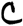 is generated with the two loci in loose linkage. Still, numerical analysis suggests that the strongest barrier is obtained at the extreme ends of the recombination scale, either for *r* → 0 or for *r* → 1 between pairs of loci. (We were not able to prove this claim, but did not find any counterexamples in numerical checks, see in SI, Fig. C24,C25.). In more detail, we find the following: First, assume that **C** appears on the island. As long as tight linkage among 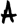 and 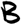 provides the strongest two-locus barrier, 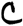 in tight linkage with 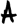 and 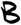 formed the strongest barrier (Fig. 5(a) and 5(b) red area, proof in section C 3.2.1). Selection against migrants is the key mechanism. If the strongest 2-locus barrier is shaped by loci 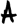 and 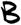 in loose linkage, we obtain the strongest 3-locus barrier for an additional adaptation 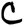 that occurs in tight linkage with either 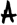 or 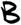. The new mutation contributes to a stronger barrier by either strengthening selection against hybrids (blue area, *γ* small, *∊_BC_* strongly negative), or by strengthening selection against migrants by reducing the direct effect of the **B** allele (green area, *∊_BC_* close to 0). Fig. 5 shows that the parameter space for having the strongest barrier in tight linkage is much larger than for loose linkage. However, genomic regions around any locus that are effectively in tight linkage are small and randomly placed loci will more likely behave as loosely linked. Therefore, optimal non-local barriers with loose linkage between 2 loci may be easier to evolve than local (island type) barriers with tight linkage among all loci.

**Figure 5:**
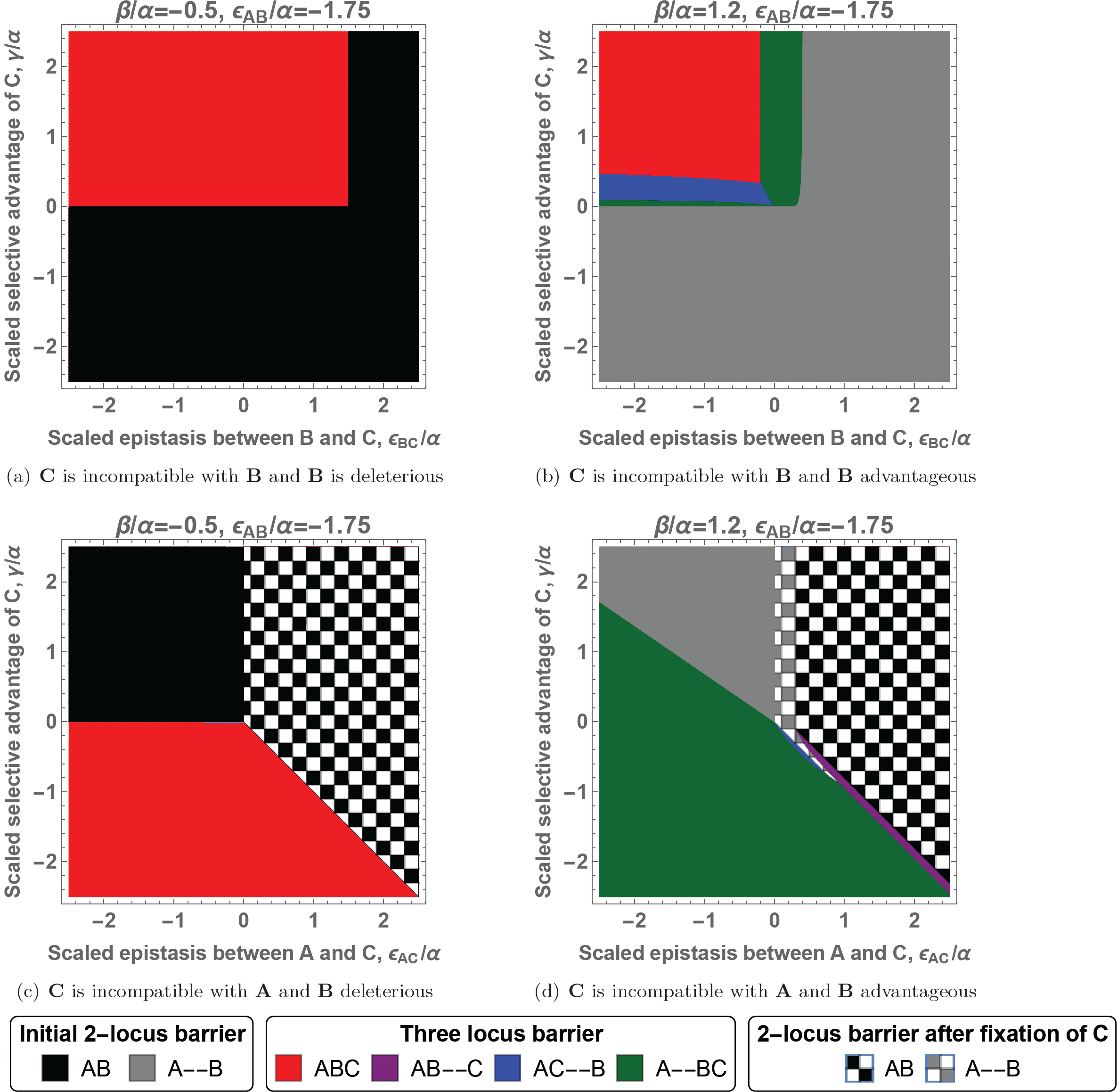
Linkage architecture forming the strongest genetic barrier for three mutations. In each panel, the x-axis corresponds to the epistasis between **C** and its interacting allele, **B** for the first row and **A** for the second row. The y-axis corresponds to the selective advantage of allele **C** on the island. The different colors indicate the linkage architecture and location a C mutation should appear to maximize 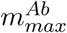. Having all loci in tight linkage can be interpreted as the existence of a single island of divergence and 2 loci in tight linkage and a third one in loose linkage as two islands of divergence, see discussion. The case of three loci in loose linkage is not represented as it never provides the strongest barrier. Analytical expressions for the barrier strengths are given in the SI, equations C7-C14. If the initial barrier remains the strongest when **C** appears on the continent, it indicates that the new mutation will always weaken the barrier. If **C** appears on the island, the barrier is unaffected by the presence of the new mutation.

If **C** appears on the continent, we observe similar results, cf. Fig 5(c) and 5(d). Having all loci in tight linkage forms the strongest barrier as long as the continental adaptations are deleterious on the island and do not generate positive epistasis (see section C 3.2.2 for proof). If having all loci in tight linkage does not generate the strongest barrier, then having 2 loci in tight linkage and the last one in loose linkage offers the strongest genetic barrier, with 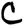 in tight linkage with 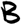, increasing both selection against migrants and hybrids (green area), or 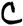 in tight linkage with 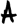, to only strengthen selection against migrants (rare, blue area). Fixing **C** is another possible mechanism to strengthen the genetic barrier if **C** generates positive epistasis with **A** (checkered areas). In this last case, the genomic location of locus ***C*** does not matter.

From these results we see that having all loci in loose linkage never seems to be the strongest linkage architecture in our model. Indeed, we did not find such an architecture despite of extensive numerical search (although we were not able to prove this). Results can be different in more complex models. After extending our model to include general 3-locus epistasis, we were able to construct a case where the strongest barrier has all three loci in loose linkage (cf. Fig. C23). However, the scenario requires a very specific type of 3-locus interaction (epistasis between **B** and **C** is only expressed in the absence of **A**) and careful fine-tuning of the selection parameters.

#### Extension of a two-locus genetic barrier, diploid populations

Here, we extend our analysis to diploid populations. More precisely, we are interested in the similarities and differences between the haploid and diploid models. The diploid model is quite complex due to the number of equations and parameters. As mentioned in the model section, we focus on two specific dominance schemes for the interactions: codominance and recessivity. Despite this simplification, only few cases (mostly when the diploid case reduces to the haploid case) allow for analytical results. In Fig. 6, we therefore compare numerical results for the strength of migration barriers.

**Figure 6:**
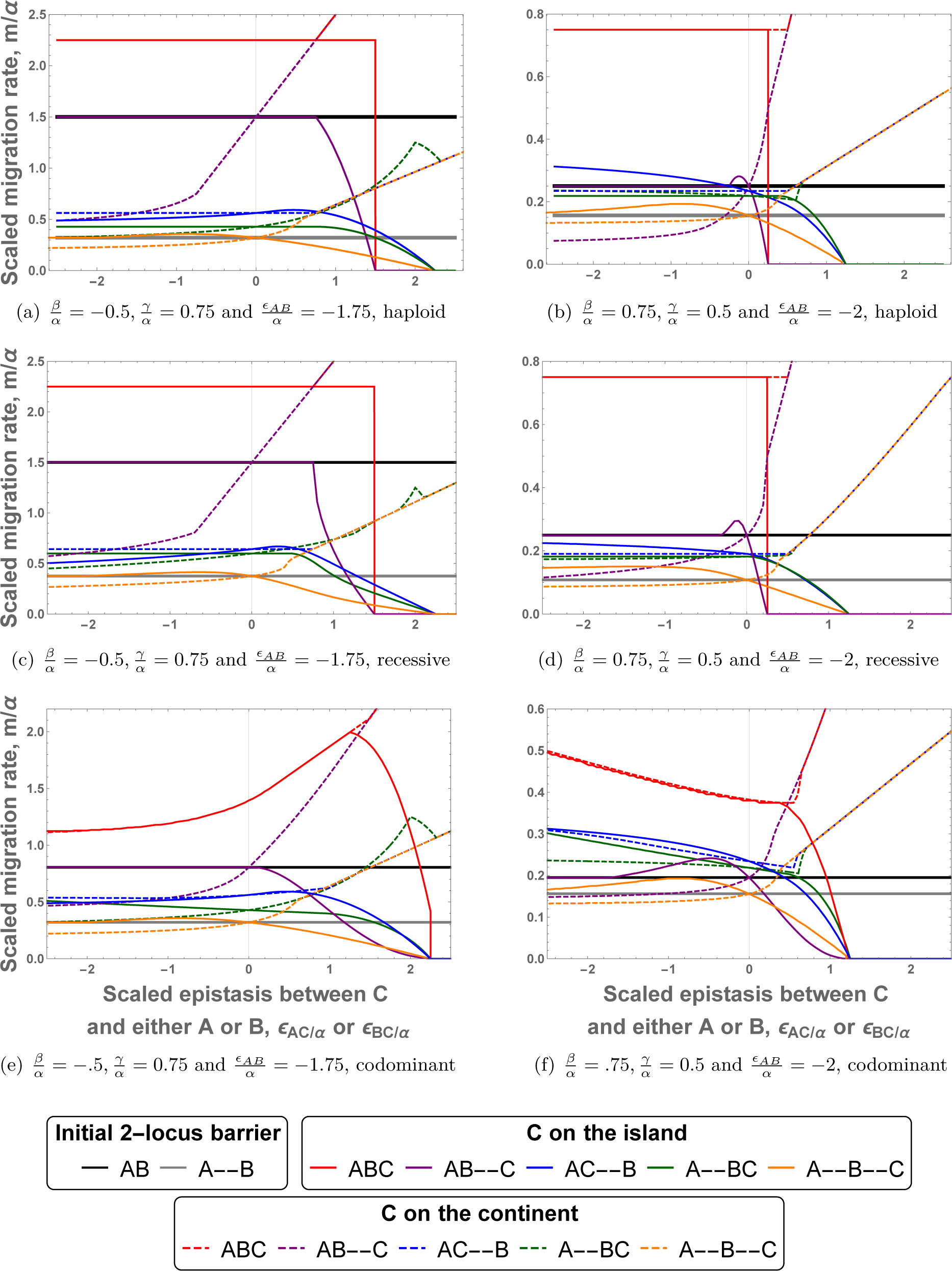
Maximal migration rate, 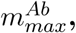 in haploid and diploid models. The X-axis corresponds to the epistatic interaction between allele **C** and its interacting allele (either **A** if **C** appears on the continent 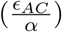 or **B** if **C** appears on the island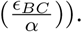 Both positive and negative epistasis are considered. The Y-axis represents the maximal migration rate for maintenance of the polymorphism at the 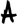 and 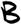 loci, 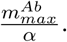 Each color corresponds to a different linkage architecture; plain lines indicate that **C** appears on the island, dashed lines on the continent. The black and grey lines serve as reference for a 2-locus genetic barrier between 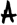 and 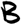 for tight linkage and loose linkage, respectively. If **C** appears on the continent, we use *−* as its selective advantage to make comparisons with **C** appearing on the island more easier (the fitness differences between the continental haplotype and the best island haplotype are then identical in the absence of epistasis). Qualitatively, the strength of the genetic barrier for given selection parameters is similar for haploid and diploid populations, with the exception of codominant epistasis between pairs of tightly linked loci.

Comparing the migration barriers for the haploid case (Fig. 6(a) and 6(b)) with recessive diploids (6(c) and 6(d)), we find broad qualitative agreement (if all loci are in tight linkage, 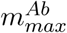 for both cases are identical). In particular, adaptation at the 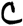 locus will weaken or strengthen the genetic barrier 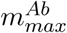 for the same linkage architectures among the three loci, and in approximately the same parameter ranges. Also having all loci in loose linkage never seems to generate the strongest barrier for a given set of parameters. Furthermore, for a given set of parameters, numerical simulations suggest that we will observe qualitatively the same optimal linkage architectures as in the haploid case when we increase *∊_BC_*.

Also the comparison of the haploid case and codominant diploids (Fig. 6(a) and 6(b) vs 6(e) and 6(f)) shows many similarities. For several architectures the dynamics (and thus the migration barriers) are identical. Indeed, as long as all interacting loci are in loose linkage, the haploid and diploid codominant model share their dynamics. This result holds for three different linkage architectures: all loci in loose linkage (orange lines) as well as 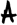 and 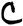 in tight linkage and 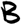 in loose linkage if **C** appears on the island (blue solid line) or 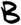 and 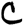 in tight linkage and 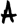 in loose linkage, if **C** appears on the continent (green dashed line). However, there is one major difference: epistasis between loci in tight linkage can be expressed. This is most noticeable when all loci are in tight linkage (red lines). Epistasis can be expressed directly in the F1 generation without any recombination event and therefore the behaviour of DMIs in tight linkage differs strongly from its haploid or recessive counterparts. Surprisingly, positive epistasis between **B** and **C**, with **C** appearing on the island, can strengthen the genetic barrier. One can relate this case to the two-locus 3-alleles model, where we have seen that reducing the hybrid cost is a viable option to strengthen the genetic barrier. A similar mechanism applies here as well.

## Discussion

How can a genetic barrier build up between two spatially separated populations that are connected by gene flow? What is the relative role of local adaptation and selection against hybrids (incompatibilities) in this process? Starting from a genetically homogeneous ancestral population, the first step of this process requires some amount of local adaptation in order to protect locally divergent alleles from swamping. For the case of (one or) two additive loci, this was discussed in detail by Bürger and Akerman (2011) and Aeschbacher and Bürger (2014), and for two loci with epistasis (allowing for incompatibilities) and unidirectional gene flow by Bank et al. (2012). Here, we have studied in more detail how a genetic barrier can be extended from such a first nucleus. As in Bank et al. (2012), we consider the case of unidirectional gene flow from a continental population to an island population.

*A priori*, there is good reason to believe that extending a barrier, once it has been initiated, should be easier than this first step (Navarro and Barton, 2003; Bank et al., 2012). Indeed, any existing divergence will reduce the effective migration rate (Barton and Bengtsson, 1986).This effect is strongest in close linkage to the first divergent locus, but also exists genome-wide; corresponding to what has been called “divergence hitch-hiking” (Via and West, 2008) and “genome hitch-hiking” (Feder et al., 2012b), respectively. It is primarily this argument that triggered the idea of islands of divergence, which may act as nuclei of emergent speciation (speciation islands, cf. Hawthorne and Via (2001); Via and West (2008); Feder and Nosil (2010); Nadeau et al. (2012); for confounding effects due to the sorting of ancestral polymorphisms see Guerrero and Hahn (2017)). If hybrid incompatibilities are involved in the build-up of such a barrier, there is a second line of argument for a subsequently increased growth of the barrier. This is the so-called “snowball effect” that predicts the accelerated growth of a genetic barrier between two allopatric populations, (Orr, 1995; Orr and Turelli, 2001), simply because with more divergent loci, there are more opportunities for incompatibilities between these loci.

Our results shed some light on the probability of observing a snowball effect in parapatry. Generally, strengthening of a genetic barrier in the presence of gene flow, while possible, is not an straightforward process, neither for haploid nor diploid populations. Indeed, although the existence of previous polymorphism can support the establishment of further divergent alleles in some cases, this process is far from being constraint-free. Furthermore, even if a new polymorphism succeeds in establishing, this does not imply that the genetic barrier is strengthened: it can as well be weakened or destroyed.

### Strengthening of an already existing genetic barrier

There are two sources from which a genetic barrier can be built or extended (Bank et al., 2012). On the one hand, selection against immigrants is effective as long as there is a fitness deficit for the immigrant haplotype relative to the island haplotype. In this case, selection acts directly to prevent introgression of the continental alleles on the island. On the other hand, selection against hybrids acts against both incompatible alleles (continent and island). It still acts as a force against introgression as long as the proportion of immigrants is small on the island. However, it is less efficient than selection against immigrants as it only acts indirectly through the selection against hybrid descendants. It also is associated with a cost for the island haplotype (production of unfit hybrids).

In the previous literature, diverse approaches have been used to study the accumulation of divergent alleles in incipient species. Flaxman et al. (2013, 2014) studied speciation with gene flow in a model without epistasis, purely through the accumulation of genes under local adaptation (selection against immigrants). As the number of locally adapted mutations increases, effective gene flow between both populations gradually declines until it reaches a so-called “congealing” threshold (sensu Turner, 1967; Barton, 1983; Kruuk et al., 1999), where effective migration rates are almost zero genome-wide and further divergence can occur at an elevated speed. The model allows for an unlimited number of local adaptation genes and generally leads to very low fitness of immigrants at (or near) speciation. There is no genetic mechanism to induce speciation in this setting: given an environment (such as a laboratory) in which both populations can survive, nothing prevents the production of viable and fertile hybrid offspring. This is at odds with theories of speciation due to the accumulation of genetic incompatibilities. Indeed, studies of allopatric speciation typically focus entirely on incompatibilities (and selection against hybrids after secondary contact) and do not include any local adaptation, e.g. Orr (1995), see also Paixão et al. (2014). Both mechanisms, selection against immigrants and against hybrids, are included in the 2-locus study by Bank et al. (2012), which we extend here.

Our results can be summarized as follows: first and foremost, we observe clear differences compared to the allopatric case concerning the accumulation of divergent alleles. Speciation in the presence of gene flow implies that each new barrier gene does not only compete against a single wildtype, but is tested against all haplotypes that can be created by gene flow and recombination. In particular, there is always selection for the reduction of hybrid cost. New adaptations on the island thus need to be locally beneficial to counter two types of costs: the direct “migration cost” to withstand swamping by the corresponding continental allele and (in case of an incompatibility) the hybrid cost. Previous divergence polymorphisms can alleviate the migration cost if a secondary adaptation occurs in close linkage, but not the cost of a stronger incompatibility. As a consequence, the number of potential barrier genes is strongly reduced relative to the allopatric case. Furthermore, there is a high probability for each new successful adaptation, on either the continent or the island, that an existing barrier will be weakened (or even destroyed) rather than strengthened. We have demonstrated this effect going from one to three barrier genes. With an increasing number of divergent genes, the constraints due to hybrid costs should only grow larger, acting against any “snowball effect”, possibly until some sort of congealing threshold (sensu Flaxman et al. (2014) Nosil et al. (2017)) is reached. Indeed, both the migration pressure and the cost of generating hybrids act as a sieve on potential new barrier genes. Due to this sieve, loci involved in DMIs (under parapatric conditions) should have on average larger direct fitness effects than loci involved in DMIs evolved in allopatry, as it has to compensate for the different costs. Furthermore, the expression of the different incompatibilities makes the process reversible as it is possible to lose some barrier genes if further adaptation reduces the hybrid cost of invading (continental) alleles. Using a model of RNA folding, Kalirad and Azevedo (2017) also found that DMIs may disappear even in an allopatric context, as further adaptations may also affect the RNA structure and therefore make the previous interactions void. Further numerical studies will be needed to quantify these predictions for general many-locus barriers.

### Migration may help to build a stronger genetic barrier to swamping

As explained above (and as expected), it is usually more difficult to extend a barrier when there is ongoing gene flow. However, we have shown that sometimes migration is necessary for a new mutation to invade. Sometimes migration can even promote adaptations (making invasion possible and/or more likely) that strengthen the genetic barrier to swamping. This can happen if the initial genetic barrier can be sustained by selection against migrants only, but the incompatibility is also strong. In that case, a new mutation generating a much weaker incompatibility, but with a weaker direct effect, can invade and strengthen the genetic barrier to gene flow if migration is strong enough. This is analogous to a reinforcement process due to pre-zygotic incompatibilities that is triggered by migration, (Kirkpatrick and Servedio, 1999). In the post-zygotic case, any adaptation that strengthens the barrier to swamping by lowering the hybrid cost (i.e. weakening the incompatibility) will make the resulting barrier more dependent on local adaptation and therefore on differences in the environment. Thus it is questionable whether this is truly a step towards reproductive isolation.

### The strongest genetic barrier for a specific set of loci

For a given set of fitness effects (both direct and epistatic) at the barrier loci, we can ask which linkage architecture provides the strongest protection against swamping. In particular: Do we get clusters of linked genes for optimal architectures?

For a two-locus barrier, Bank et al. (2012) have shown that the most stable architecture is always one with extreme linkage. If the barrier is primarily maintained due to selection against immigrants, tight linkage (*r* = 0) results in the strongest barrier (this is always the case in the absence of epistasis (Akerman and Bürger, 2014)). In contrast, the most stable barrier is obtained with maximally loose linkage (corresponding to *r → 1* in the model) if selection mainly acts against hybrid recombinants.

When we extend the 2-locus model to three loci, we can distinguish three patterns with extreme linkage among pairs of loci: all three loci in tight linkage, two tightly linked loci and one loosely linked locus, or all three loci loosely linked. We find examples, for each of three patterns mentioned above, where the considered linkage architecture formed the strongest genetic barrier.. However, the pattern of three loosely linked loci seems to be very rare. We only observe this result in a custom-made model with 3-way epistasis among the three loci and restricted to a small parameter range. Usually, we obtain either one or two islands of divergence: one if continental adaptations are deleterious on the island and two otherwise. As in the 2-locus case, we do not observe (based on a limited number of numerical studies) an optimal architecture involving intermediate recombination, even if the genetic barrier is no longer a monotonic function of recombination and local maxima of the barrier strength as function of the recombination rate exist in some cases (Fig. C24(b), blue dashed line). Note however that such an optimum at intermediate recombination distances can occur in stochastic models (Aeschbacher and Bürger, 2014).

Although stable architectures will be favored in the presence of gene flow, the most stable barriers are not necessarily the ones that will evolve most easily in natural populations. For most selection parameters, two or more loci in tight linkage provide the strongest barrier. However, the area around each single locus that behaves as essentially tightly linked is usually very small relative to the size of the genome. Thus, if interacting genes are scattered across random positions in the genome, stable configurations will be rare. Chromosomal rearrangements such as inversions can procure larger regions of no recombination, increasing the likelihood of barrier loci in tight linkage. Navarro and Barton (2003) discussed the importance of such rearrangements in the speciation process. However, gene conversion can also occur in inversions (Korunes and Noor, 2017). In addition, a study between two *Senecio* species (Brennan et al., 2014) found no associations between those rearrangements and incompatible genes.

New adaptations that appears at loosely linked loci could be transient. As demonstrated by Yeaman (2013), adaptations that first occur in different genomic regions, can later move into tight linkage due to genome rearrangement. Indeed, some studies, reviewed in Feder et al. (2012a), report small regions of divergence hitch-hiking in several species, suggesting that there may be at least a weak trend for an accumulation of divergent sites. Currently, however, neither theory nor empirical evidence provide a strong basis for divergence islands as a reliable pattern for parapatric speciation.

### Biological evidence and implications

Growth of a genetic barrier starting from an initial pairwise DMI could be common in nature. Indeed, Corbett-Detig et al. (2013) reported that two locus DMIs already exist within populations of the same species, with an average of 1.15 DMIs between different *Drosophila melanogaster* recombinant inbred lines that were derived from a common parental pool. Segregating incompatibilities have also been found for yeast (Marsit et al., 2017). This suggests that speciation through the accumulation of post-zygotic incompatibilities may not start from scratch (a common hypothesis in many models), but can rely on divergence that already exists between populations. This makes the process investigated here (how new mutations can strengthen a genetic barrier) a crucial step of the speciation process.

Dettman et al. (2008) evolved populations of *Neurospora* in two different environments. They crossed individuals that had evolved independently (in allopatry) either in the same environment (parallel evolution) or in a different environment (divergent evolution). Since crosses between individuals under different selective pressure tended to generate more unfit individuals, they concluded that genes involved in early divergence also generate genetic incompatibilities, see also Kulmuni and Westram (2017). This corresponds to the assumptions of our model. In addition, in our model we do not consider independent DMIs but partially saturated ones (one locus involved in two DMIs). Guerrero et al. (2017) provided an example of such saturated incompatibilities in the pollen of two species of *Solanum* genus.

Ono et al. (2017) measured the epistasis between first-step adaptations within a pathway responsible for fungicide resistance in yeast, with each mutation in a different gene. They found pervasive epistasis among these mutations, with a third of the interactions classified as DMIs. Based on these findings, they suggested a scenario of parallel adaptation (without local adaptation) for allopatric speciation with secondary contact: if populations that adapt in parallel to the same environment (during an allopatric phase) fix different mutations in the same pathway, incompatibilities can easily be generated. Our model predicts that this kind of adaptation (*α* ≈ *β*, i.e. only selection against hybrids) can only be maintained in the face of gene flow when the DMI loci are far away from each other. This is indeed the case for Ono et al. (2017): the mutations involved in DMIs occur in genes located on different chromosomes. In addition, our model (and Bank et al. (2012)) shows that an allopatric phase is not needed for the evolution of such a DMI: it can also evolve with ongoing unidirectional gene flow given that the first substitution happens on the island. For bidirectional gene flow, local adaptation is required. Note that when extended to diploids, the constraint on the linkage architecture vanishes if the incompatibility is expressed in F1 hybrids (codominant incompatibility).

### Model assumptions and possible extensions

Our model relies on a number of hypotheses, most of them shared with Bank et al. (2012). First, we assume an infinite population size to ignore the effects of genetic drift. This assumption is adequate as long as the population size is large enough that drift can be ignored relative to the other evolutionary forces 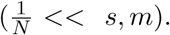 We only study whether the mutant can invade or not, but do not consider establishment probabilities. When these are included (Aeschbacher and Bürger (2014) for two loci without epistasis), the highest establishment probability is often not found for the most stable configuration (tight linkage in this case), but for small, non-zero recombination rates.

We focus entirely on a continent-island model with unidirectional migration. This is realistic if either physical mechanisms enforce unidirectional gene flow (wind, water current, flowering time), or if the contribution of both populations to a common migrant pool is strongly biased (because of unequal population size or because of reduced fertility, e.g. of a marginal population). If there is weak back migration, the effects described here should still hold, as long as the island adaptations are not advantageous on the continent. For strong bi-directional migration, generalist genotypes can gain an advantage and different results are obtained (Akerman and Bürger, 2014).

Due to the complexity of the system, we restrict our analytical analysis to the limiting cases of recombination (*r* = 0 and *r* = ∞). Bank et al. (2012) have shown that the analysis of these limiting cases provides a good understanding of the general case for the 2-locus model. Our numerical study for intermediate recombination rates confirms this for three loci, Fig. C24, C25. We assume that all loci are autosomal. Höllinger and Hermisson (2017) provide an analysis of a two-locus DMI in parapatry for organelles and sex chromosomes.

Finally, we have restricted our detailed analysis in this paper to epistasis schemes with only pairwise interactions. Complex epistasis networks with interactions linking three or more loci offer further routes to strengthen a genetic barrier that will be explored in a forthcoming study.

## Acknowledgments

We thank R. Bürger, M. Servedio, C. Vogl, S. Mousset, C. Bank, I. Fragata, I. Höllinger and the Biomathematics Group at the University of Vienna for helpful discussion and comments on the manuscript. We thank the two anonymous reviewers for their valuable suggestions that have improved this manuscript. AB was supported by the Marie Curie Initial Training Network INTERCROSSING.

